# Human Cytomegalovirus Induces Significant Structural and Functional Changes in Terminally Differentiated Human Cortical Neurons

**DOI:** 10.1101/2023.03.03.531045

**Authors:** Jacob W. Adelman, Suzette Rosas-Rogers, Megan L. Schumacher, Rebekah L. Mokry, Scott S. Terhune, Allison D. Ebert

## Abstract

Human cytomegalovirus (HCMV) is a highly prevalent viral pathogen that typically presents asymptomatically in healthy individuals despite lifelong latency. However, in 10-15% of congenital cases, this beta-herpesvirus demonstrates direct effects on the central nervous system, including microcephaly, cognitive/learning delays, and hearing deficits. HCMV has been widely shown to infect neural progenitor cells, but the permissiveness of fully differentiated neurons to HCMV is controversial and chronically understudied, despite potential associations between HCMV infection with neurodegenerative conditions. Using a model system representative of the human forebrain, we demonstrate that induced pluripotent stem cell (iPSC)-derived, excitatory glutamatergic and inhibitory GABAergic neurons are fully permissive to HCMV, demonstrating complete viral replication, competent virion production, and spread within the culture. Interestingly, while cell proliferation was not induced in these post-mitotic neurons, HCMV did increase expression of proliferative markers Ki67 and PCNA suggesting alterations in cell cycle machinery. These finding are consistent with previous HCMV-mediated changes in various cell types and implicate the virus’ ability to alter proliferative pathways to promote virion production. HCMV also induces significant structural changes in forebrain neurons, such as the formation of syncytia and retraction of neurites. Finally, we demonstrate that HCMV disrupts calcium signaling and decreases neurotransmission, with action potential generation effectively silenced after 15 days post infection. Taken together, our data highlight the potential for forebrain neurons to be permissive to HCMV infection in the CNS, which could have significant implications on overall brain health and function.

## INTRODUCTION

Human cytomegalovirus (HCMV) is a common pathogen with a worldwide infection rate ranging from 60 to 90% of the population (1–3). Though infected individuals are typically asymptomatic, a subset of congenital infections (10-15%) demonstrate a risk to fetal health and development (4–9). In such cases, a wide range of neurological symptoms may occur, including hearing impairment, cognitive deficits, learning/language disorders, and microcephaly (2, 9–11). Interestingly, several studies have also examined neurological effects present in individuals with asymptomatic congenital infections (2, 4, 12, 13). Together, these studies highlight HCMV’s ability to negatively impact development and activity within the central nervous system (CNS).

The role of HCMV within the CNS is likely complex and multifaceted. Previous reports demonstrate that HCMV can infect several resident cell types within the CNS, including astrocytes (14–16), oligodendrocytes (17), and neural progenitor cells (NPCs) (7, 18–21). However, the ability of neurons to act as a susceptible target of HCMV infection is often debated, with empirical results ranging from no detected infectivity to full permissiveness (14, 20–25). Among those demonstrating a lack of HCMV infection within neurons, the cells’ post-mitotic state is commonly indicated to be the underlying reason (14). Functionally, little is known about the effects of HCMV on terminally differentiated neuronal populations, though early reports detail tendencies toward cell death and altered responses to glutamate stimulation (25). This effect is in addition to functional aberrancies observed in other cell types such as NPCs and fibroblasts, often involving dysregulation of calcium signaling (26–28). Considering the limitations of existing data, further analysis of the effect of direct infection on neurons is warranted.

Induced pluripotent stem cells (iPSCs) represent an indispensable tool for studying effects on human tissues. This is evident through both their robust differentiation capabilities – allowing for the study of viral effects on various cell types – and their species compatibility with HCMV. To date, a majority of studies analyzing the effect of HCMV infection on CNS cells have been conducted using either fetal-derived primary cultures or cell type-specific lines (7, 14, 15, 17, 19, 20, 29, 30), while, in contrast, few have utilized iPSC-derived cultures (25, 26, 31). Further, iPSCs allow for convenient, accurate modeling of neuronal populations present in the human forebrain; an area that is both responsible for higher cognitive functions and influenced by congenital HCMV’s neurological symptoms. The iPSC-derived neurons may be particularly relevant for studying the role of HCMV in neurodegenerative conditions such as Alzheimer’s disease (AD) (32–34).

In the current study, the structural and functional effects of HCMV infection were studied in the context of iPSC-derived, terminally differentiated forebrain neurons. We assessed HCMV replication from immediate-early gene expression through virion production in terminally differentiated and functionally mature iPSC-derived excitatory and inhibitory neurons. We discovered that these mature neurons are highly permissive to HCMV infection, exhibited viral replication based on expression of the late protein pp28, and can produce infectious virus. Finally, we show that HCMV infection in terminally differentiated neurons essentially eliminated action potential generation and calcium signaling. These results demonstrate that terminally differentiated human neurons are permissive to HCMV and that infection significantly alters both structural and functional features of mature neurons. As such, these data suggest that HCMV infection could have pathological consequences in the adult CNS.

## RESULTS

### Terminally differentiated forebrain neurons derived from iPSCs support HCMV lytic replication

Utilizing iPSCs from five unrelated adult individuals (3 healthy control lines, an AD patient line with a PSEN2_N141I_ mutation, and an AD patient with sporadic AD (sAD)), forebrain neurons were differentiated through a neural progenitor cell (NPC) intermediate (Fig. 1A). As HCMV presence has been documented to inhibit neural differentiation (7, 19), cultures were propagated for 84-150 days of differentiation (assay dependent) prior to infection to ensure presence of functionally mature neurons in culture. Neurons examined at Day 84 of differentiation show the dominant presence of both inhibitory GABA-ergic (GABA^+^) neurons and excitatory glutamatergic (VGLUT2^+^) neurons (**Fig. S1A**). Additionally, vimentin^+^ astrocytes a can be found within the cultures (**Fig. S1A**). Consistent with neuron maturation, we observe a change in cell morphology over time with NPCs exhibiting a more cobblestone appearance and lacking defined neurite projections to greater neurite elongation, development, and network establishment in late cultures (**Fig. 1A and 2**). As expected, we observe a reduction in expression of the NPC marker Pax6 and a near complete loss of Ki67 expression over time (**Fig. 2**). To establish that the observed infection phenotypes are not an artifact of the iPSC-derived neuronal differentiation paradigm, we conducted additional experiments using neurons generated from the fetal cortex. Fetal cells were cultured for ∼20 weeks as nonadherent neurospheres (35, 36) and then plated as a monolayer and differentiated for an additional 30 days. We found GABA^+^ and VLGUT2^+^ neurons, robust neurite outgrowth, lack of Pax6 expression, and minimal Ki67 staining, consistent with the iPSC-derived neurons (**Fig. S1B and 2**). Together, these data establish the iPSC model system and clearly demonstrate the presence of post-mitotic differentiated neurons.

**FIG 1.**
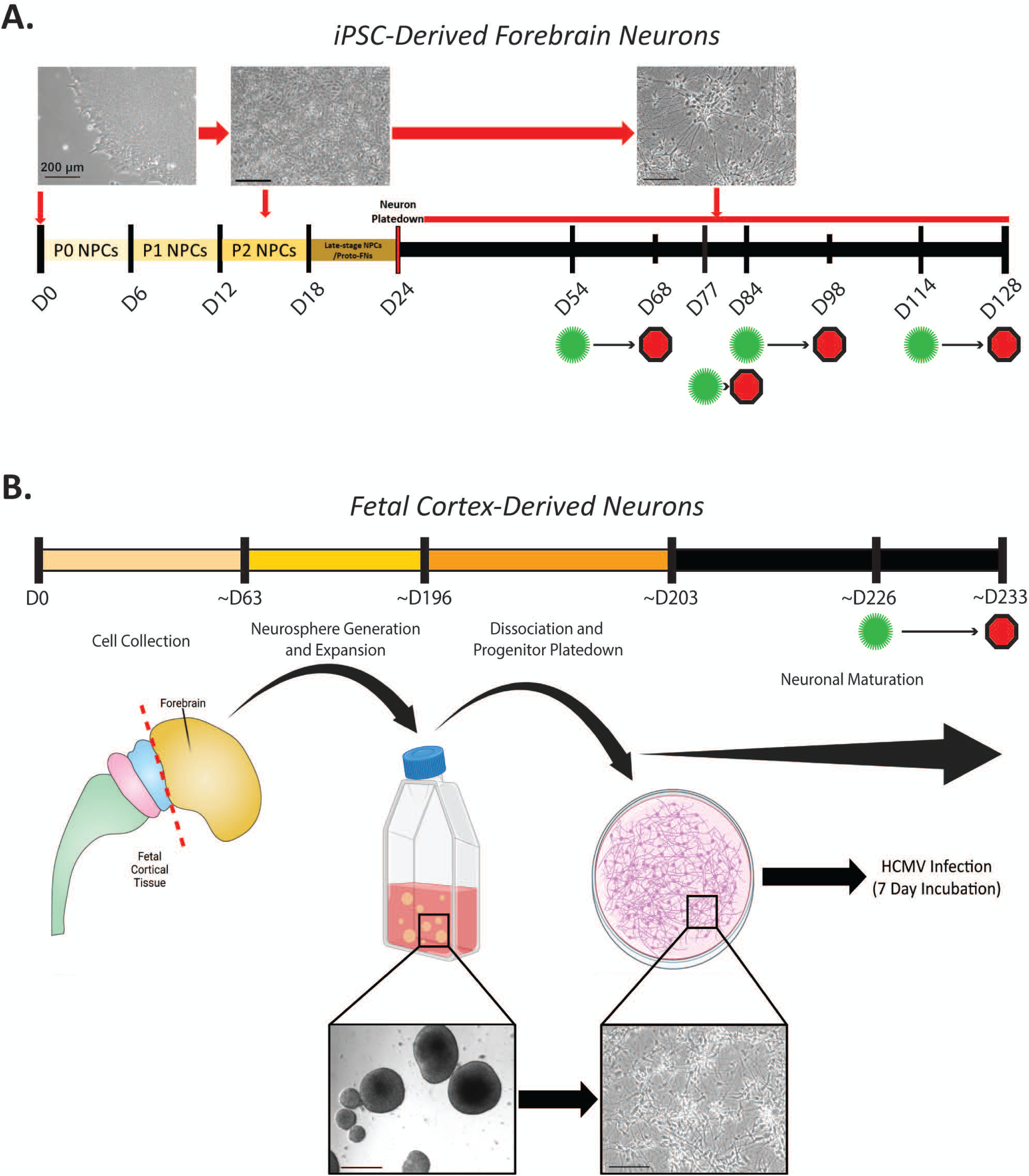
Assessed neurons were generated from both iPSC- and fetal-derived progenitor cells. (A) iPSC colonies were dissociated, plated into SMADi Neural Progenitor Medium, and differentiated into neural progenitor cells over an 18-day period. Generated NPCs were dissociated/plated into forebrain media differentiation medium for 6 days to prepare immature forebrain neurons. Final plating of immature neurons occurred at D24 of differentiation and cells were maintained in forebrain neuron maturation medium until use. Time points of HCMV infection are denoted by green virus icons and culture collections are indicated by red octagons. (B) Fetal-derived cultures were collected from forebrain tissue at ∼D63 of in-utero development (∼9 weeks). Neural progenitor samples were cultured as neurospheres and expanded for an average of 133 days (19 weeks) prior to neuronal differentiation. Neurospheres were differentiated over a 1-week period and plated down at ∼D203 of differentiation. Plated neurons were allowed to grow for 23 days prior to HCMV exposure (D226), with infection persisting for an additional 7 days (D233) prior to sample collection.

**FIG 2.**
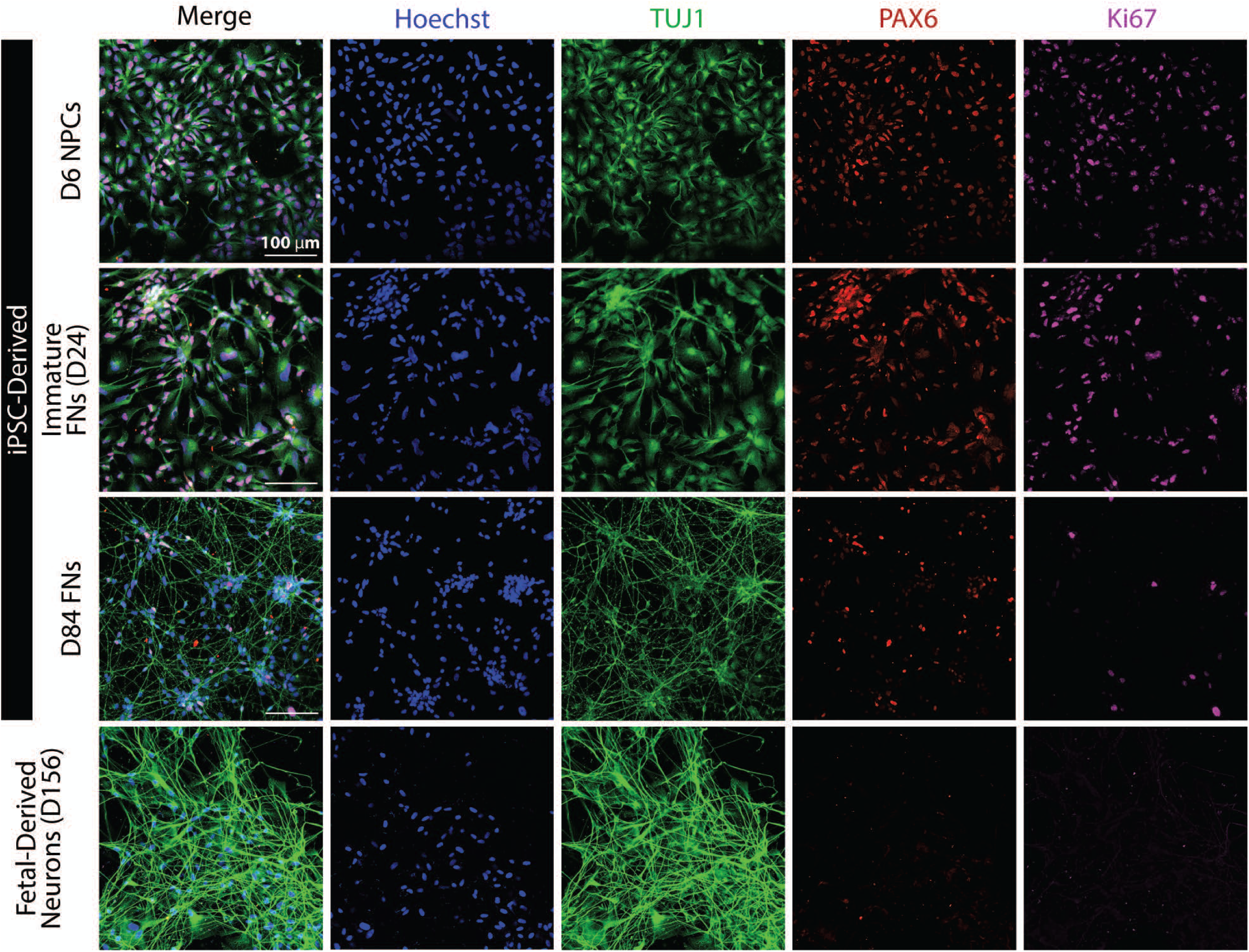
iPSC- and fetal-derived cells demonstrate appropriate neuronal morphology and molecular signatures. In iPSC-derived cultures, neuron-specific class III beta-tubulin (TUJ1) staining is refined into distinct cell processes as cultures mature from NPCs to forebrain neurons. Additionally, NPC marker Pax6 and proliferative marker Ki67 are decreased throughout differentiation and maturation, indicating that neurons have become postmitotic. These features are recapitulated in the fetal-derived neurons, with cell demonstrating clear TUJ1^+^ processes and minimal Pax6 and Ki67 positivity.

With the culture system established, we next infected D30 forebrain neurons with HCMV originating from BAC cloned TB40/E strain. We used two recombinant versions, HCMV TB40/E-eGFP, containing the EGFP gene under control of an SV40 promoter (37, 38), and TB40/E-eGFP/mCh expressing late protein pp28 in-frame with the fluorescent protein mCherry and IE2 in-frame with a cleavable eGFP (IE2-2A-eGFP UL99-mCh) (39). We infected mature neutrons at 0.5, 1, and 3 IU/cell (**Fig. 3A**). Virus was added to forebrain neurons for 2 hrs, after which the inoculum was removed, and fresh medium added every other day for the duration of the experiment. For both viruses, we observed fluorescence at all MOIs within 48 hours post infection (hpi) with an MOI of 3 being most robust (**Fig. S2A**). By 15 days post infection (dpi), differences in fluorescence between MOIs appear negligible (**Fig. S2B**). Expression of pp28-mCherry in forebrain neurons cultures confirmed that infection was proceeding from the initial step of viral entry through late gene expression (**Fig. 3B and 4A**). Further, to confirm infection in a second model system, fetal cortex-derived neurons were infected with TB40/E-eGFP and demonstrated a progressive increase in the number of GFP^+^ cells over time (**Fig. S3**).

**FIG 3.**
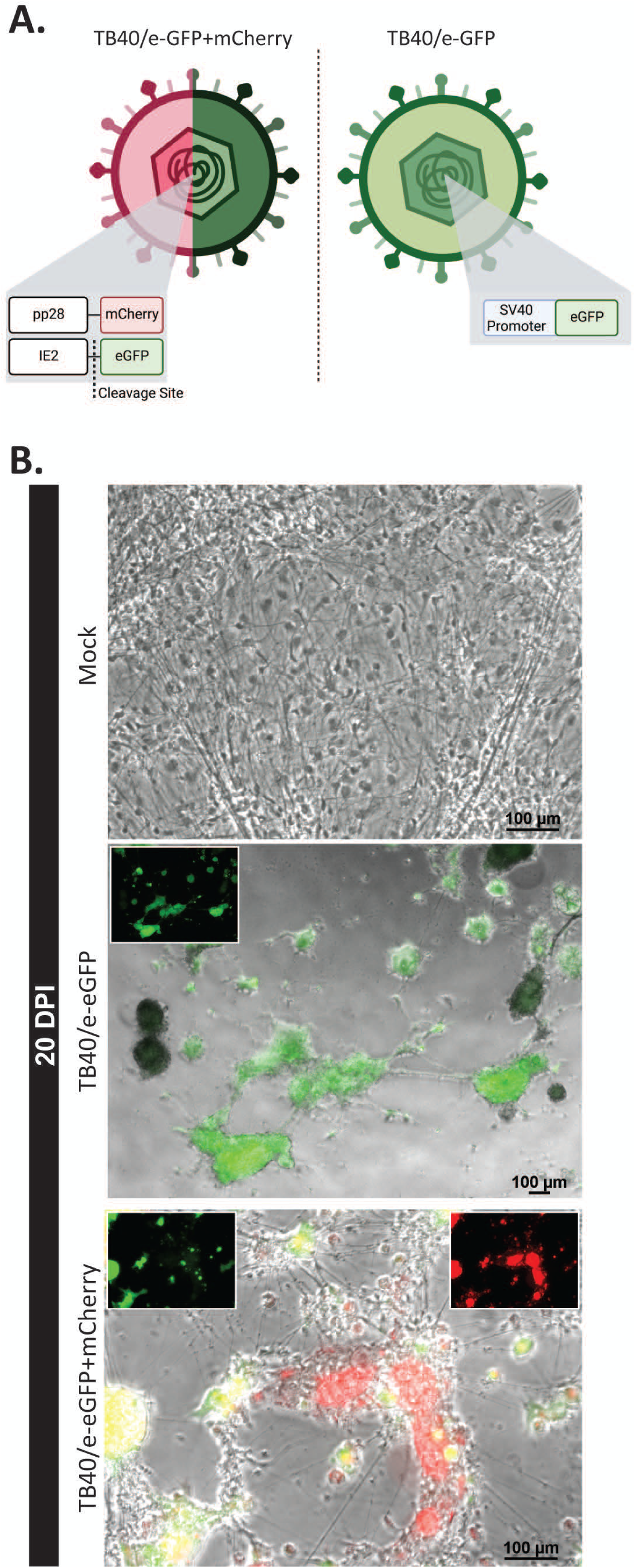
HCMV clinical strain TB40/e subvariants can infect and replicate within forebrain neuron cultures. (A) TB40/e-GFP+mCherry contains an eGFP marker of immediate early gene (IE2) synthesis and an mCherry fluorophore linked to late gene product pp28. TB40/e-GFP utilizes an SV40 promoter-driven eGFP to denote infection. (B) At 20 dpi, forebrain neurons cultures infected with both viral constructs demonstrate robust fluorescence.

**FIG 4.**
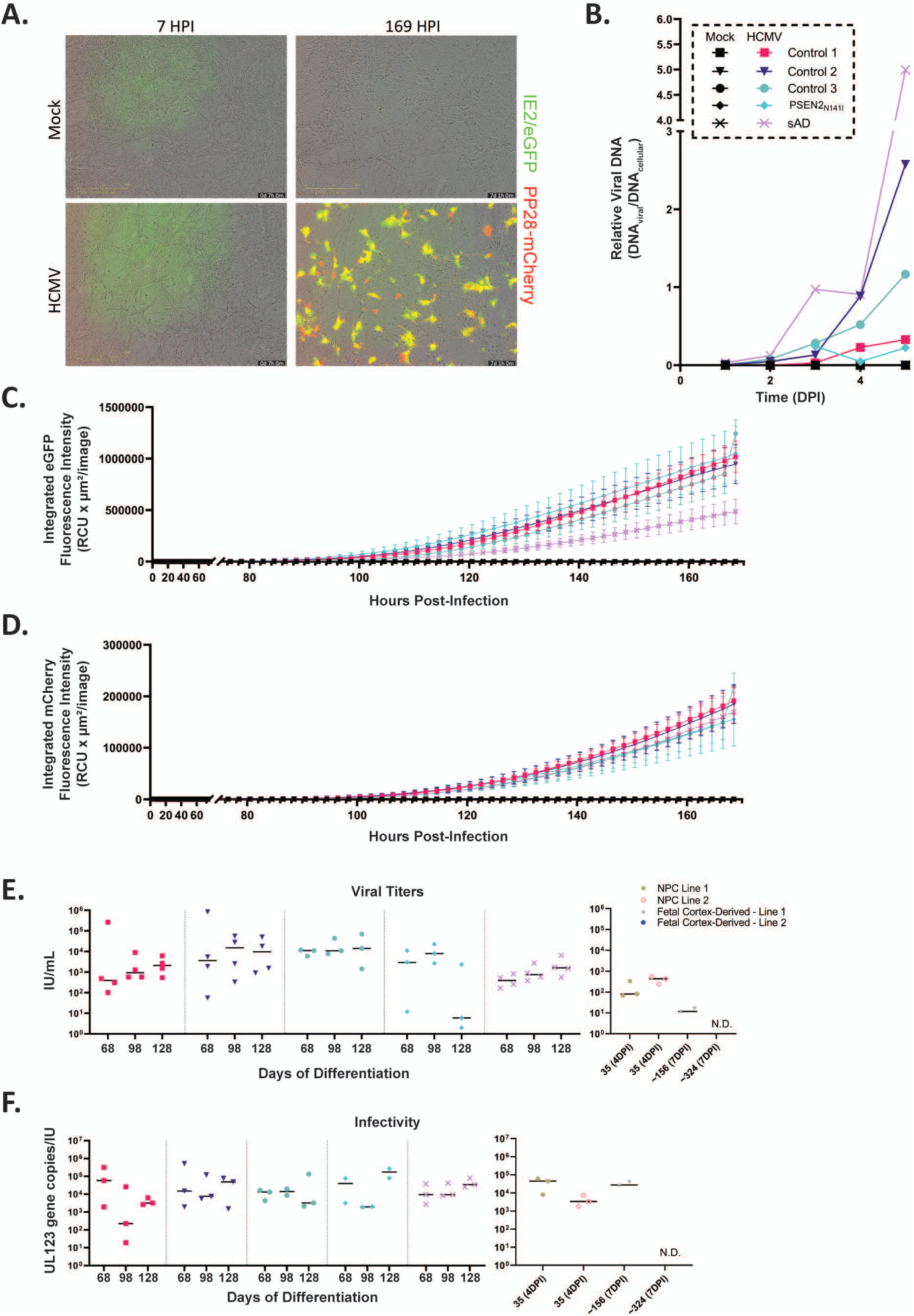
HCMV infection is progressive and capable of producing functional virus within neuronal cultures derived from iPSC and fetal progenitors. (A) Live imaging reveals the increase in both eGFP and mCherry signal between 7 hpi and 169 hpi. (B) Analysis of relative viral-to-cellular DNA details progressive increases in viral burden within forebrain neuron cultures over the first 5 days of infection (n = 1-3 biological replicates). (C-D) Supporting increases in viral DNA expression, fluorescence measurements highlight increases in both eGFP and mCherry throughout the initial 7 days after HCMV exposure (n = 3). (E) Assessment of viral titers within 14 dpi neuron conditioned media (inoculum) was completed using ARPE-19 epithelial cells. All tested forebrain neurons cultures produced relatively consistent viral titers regardless of the timepoint being assessed. These values are comparable to those seen in tested neural progenitors at 4 dpi. Fetal-derived neurons (7 dpi) demonstrated inconsistent results, with one of two lines producing viral titers (n = 2-4). (F) Infectivity of viral products was assessed using copies of UL123 present within neuron conditioned media relative to that sample’s viral titer, shown in E. As with the titer values, infectivity was consistent across all forebrain neuron lines and timepoints. Further, forebrain neurons values were consistent with NPC infectivity ratios. As in E, fetal derived cells produced highly variable infectivity.

To evaluate viral spread in long-term forebrain neuron cultures, time course data were collected tracking infection from point of infection (D53 of differentiation) through 7 dpi. Robust eGFP and mCherry signals were detected over the duration of the experiment, with some cells demonstrating primarily late gene (pp28) expression by 7 dpi (**Fig. 4A**). In conjunction with imaging experiments, forebrain neuron cell pellets were collected at daily timepoints, processed via DNA extraction, and evaluated for relative viral-to-cellular DNA over time. Across each line, relative viral DNA load increased over 5 dpi (**Fig. 4B**). Supporting these data, fluorescence tracking revealed the expression of early gene proteins (observable via eGFP) becoming prominent around 90 hpi. These results were notably consistent between the three controls and the PSEN2_N141I_ line, whereas the sAD line demonstrated a lower amount of eGFP fluorescence. Late gene expression (mCherry) was noted shortly thereafter, with fluorescence increasing near 100 hpi (**Fig. 4C and D**). All lines were consistent with regard to amount and rate of mCherry fluorescence over time.

After confirming that forebrain neurons could be infected by HCMV, we next asked if infected forebrain neurons could produce functional, infectious virus. At D84, D114, and D144 of differentiation, cells were infected with TB40/E-eGFP for 2hrs, washed, and cultured in fresh medium for 14 days. Conditioned media from forebrain neurons were collected and applied to ARPE-19 epithelial cells to determine viral titers. Across all timepoints, cells from each of the iPSC lines produced infectious virus (**Fig. 4E**). Further, no significant differences were found between cell lines at each timepoint. It should be noted that although viral titers were lower than those expected from epithelial cells and fibroblasts, the production of functional virus in neural cultures signifies viral permissiveness (40). iPSC-derived neurons were also compared to both iPSC-derived NPCs and fetal cortex-derived neurons. iPSC-derived neurons demonstrated little difference as compared to iPSC-derived NPCs (**Fig. 4E**). Fetal-derived neurons were less consistent, with one of the tested lines showing a slight reduction in viral titer and the other no viral titer (**Fig. 4E**). To further examine virus production, viral infectivity of the resulting virus was assessed across all timepoints using the ratio of UL123 gene copies per infectious units. Again, across all timepoints and lines, average infectivity ratios were relatively similar (**Fig. 4F**). Interestingly, forebrain neuron cultures were determined to produce larger amounts of non-functional virus relative to other infected MRC-5 fibroblasts (41). Comparisons to NPCs and the permissive fetal-derived culture revealed no significant differences compared to forebrain neuron cultures (**Fig. 4F**). Taken together, these data demonstrate that iPSC-derived forebrain neurons are susceptible to HCMV infection and support the full viral replication cycle.

### Infection by HCMV stimulates cell cycling markers in post-mitotic forebrain neurons

Infection by HCMV manipulates cell cycle in human fibroblasts and salivary gland epithelial cells (42–45). Mature neurons are post-mitotic, having entered G0 phase upon full differentiation from NPCs (46). As such, the effects of HCMV on the neuronal cell cycle are largely unknown. To determine if HCMV infection reverts differentiated neurons back into a mitotic state, we evaluated expression of several cell cycle proteins. Western blot analysis of D98 forebrain neurons (14 dpi) demonstrated HCMV-dependent decrease in the G2-to-M Cyclin B in all lines (**Fig. 5A and 5B**). We then analyzed the protein expression of cyclin-dependent kinase inhibitor p21 and infection did not appear to have any consistent effect on its expression (**Fig. 5C**). In contrast to Cyclin B and p21, infection resulted in a consistent increase in PCNA expression. PCNA is an essential protein in DNA synthesis is often elevated in S phase. We next assessed expression of a marker of proliferation, Ki67. Using immunofluorescence in D84 (7 dpi) cultures, significant increases in Ki67 staining were detected within three of the four tested forebrain neuron cultures. Despite increased PCNA and Ki67, we did not observe a subsequent increase in the cell density (**Fig. 5D-E**). These results are in line with previous studies using multiple cell lines and HCMV strains (47–49). Furthermore, Ki67 staining colocalized with areas of infected cells (**Fig. 5D**). Taken together, these data demonstrate that HCMV infection of forebrain neurons stimulates markers of proliferation consistent with events occurring during infection of non-postmitotic cell lines.

**FIG 5.**
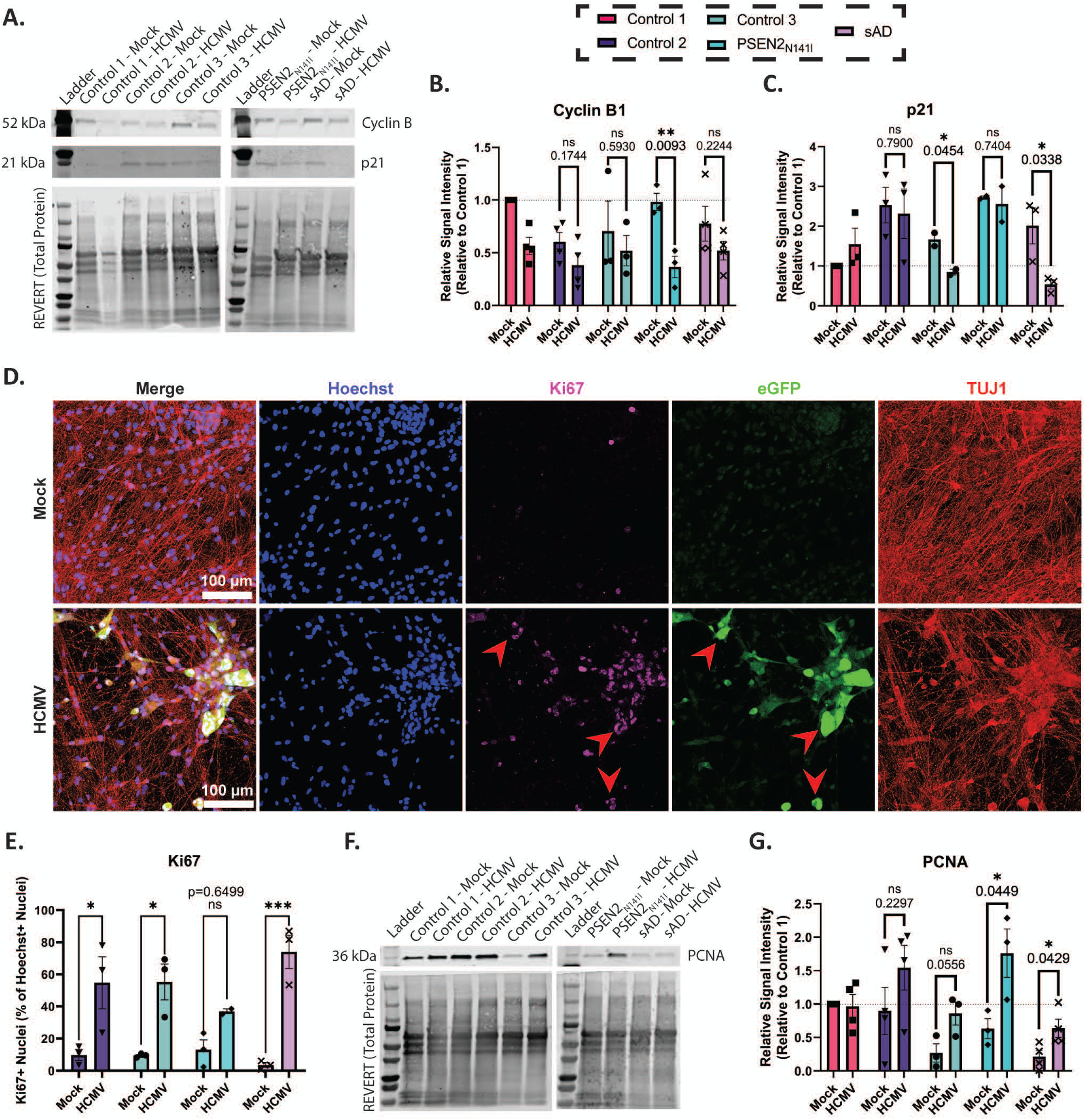
HCMV infection alters key targets associated with cell cycling. (A) Representative western blot showing banding patterns for cyclin B1 and cyclin-dependent kinase inhibitor 1 (p21), with included total protein stain. (B) Quantification of cyclin B1 western blots demonstrates a trend of HCMV downregulation across all lines, with significance in the PSEN2N141I forebrain neurons (t-tests. (C) Analysis of p21 blotting reveals an unclear trend, with two of the five tested lines demonstrating significant downregulation. (D) Representative immunofluorescent staining for proliferative marker Ki67. Key areas of overlap between Ki67 and viral eGFP are denoted with red arrows. (E) In both controls and the sAD line, HCMV induces an increase in Ki67+ cells. (F) Representative Western Blot demonstrating banding patter for proliferative cell nuclear antigen (PCNA), with included total protein stain. (G) Quantification of PCNA blotting demonstrates a trend of HCMV-mediated upregulation in forebrain neurons. Data are presented as mean ± SEM, with n = 2-4. Student’s t-tests were used to determine statistical differences in B, C and G, while two-way ANOVA was used in E. *, p < 0.05; **, p < 0.01; ***, p < 0.001

### Infection alters human forebrain neuron structure

We and others have noted an increase in apoptosis within human NPCs and cerebral organoids upon HCMV infection (19, 50, 51). Therefore, we next tested whether HCMV induced cell death of terminally differentiated forebrain neurons. We used terminal deoxynucleotidyl transferase dUTP nick end labeling (TUNEL) and found no significant increase in TUNEL positive cells in the HCMV infected cultures to mock controls at D84 (7 dpi) (**Fig. 6A and B**).

**FIG 6.**
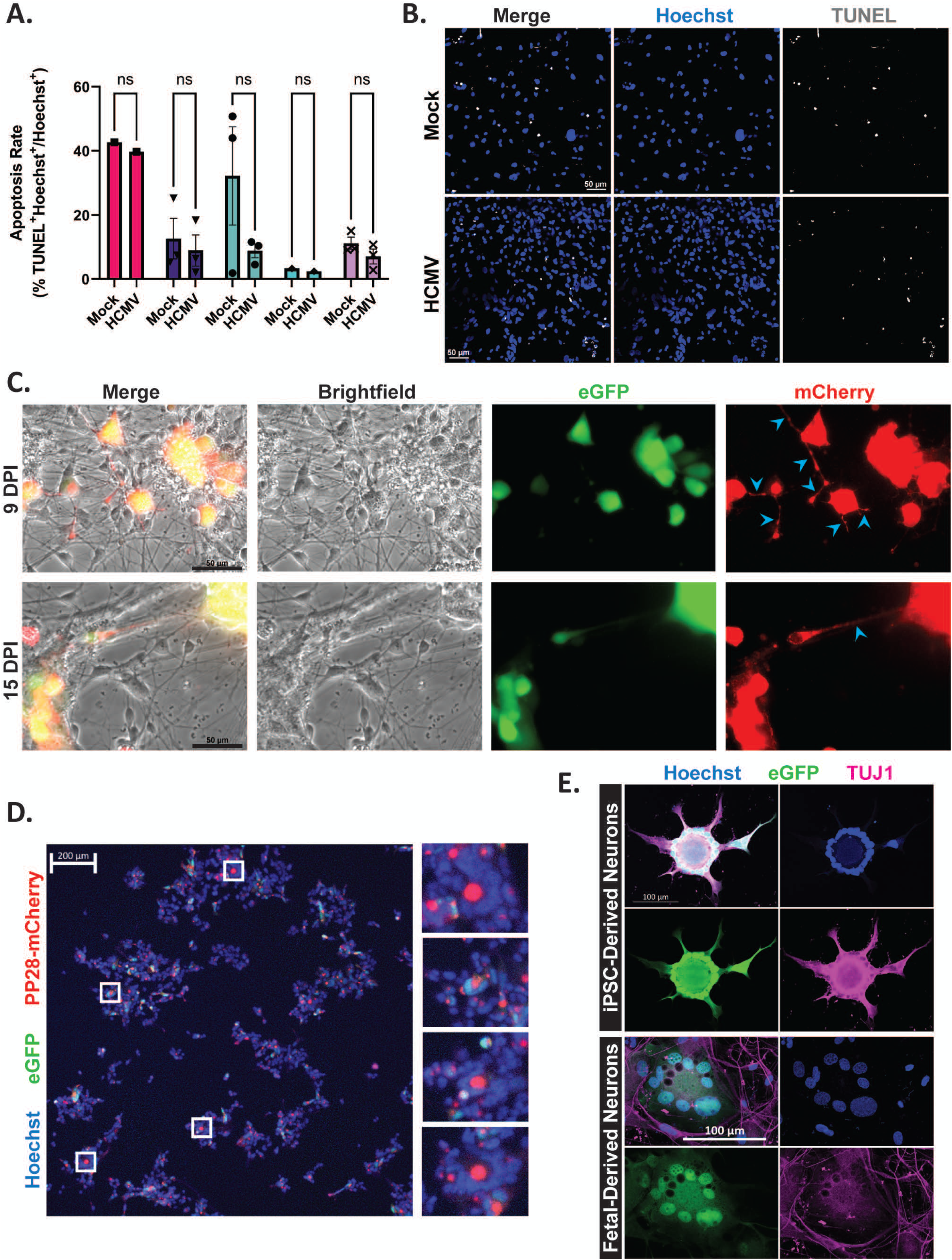
HCMV infection significantly alters neuronal structure while maintaining cell viability in forebrain neurons. (A) Assessment of nuclei for the presence of TUNEL signal revealed that, while apoptotic rates varied by cell line, there was no significant difference between HCMV infected cultures and mock-treated cells. (B) Representative image of TUNEL staining in mock- and HCMV-treated Forebrain neurons. (C) Infection of forebrain neurons with TB40/e-GFP+mCherry revealed the presence of pp28-mCherry signal within neuronal processes, indicating a potential trafficking mechanism (indicated with blue arrows). (D) Using TB40/e-GFP+mCherry, pp28 signal can also be observed organizing into virion assembly compartments (VACs) surrounded by rings of nuclei (syncytia). (E) Syncytial structures are formed in both iPSC-and fetal-derived cultures. Further, TUJ1 positivity indicates that neurons are the components of the observed syncytia.

Neuronal infection with the alpha-herpesvirus herpes simplex virus (HSV) has been shown to be transported from the cell body through neurites (45, 52–54). Therefore, we next examined pp28-mCherry localization in both the cell body and in neurite projections. We infected D84 forebrain neurons with TB40/E-eGFP/mCh and observed pp28-mCherry expression within neuronal projections and large accumulations of pp28-mCherry within cell bodies at 9 and 15 dpi (**Fig. 6C**). pp28 is proposed to be trafficked to the cytoplasmic Virion Assembly Compartment (VAC), a process necessary for virion assembly (55). We observed VAC formation in forebrain neuron cultures indicated by large, punctate regions of pp28 (**Fig. 6D**).

Syncytia formation occurs during HCMV infection whereby disparate cells fuse to become a single, multinucleated cell (56–58). We observed forebrain neurons infected with HCMV do fuse into syncytia-like formations as characterized by a pp28-positive VAC surrounded by a ring of nuclei (**Fig. 6D**). These structures were also present in cultures infected with the TB40/E-eGFP (**Fig. 6E**). To determine if the cells composing the syncytia were once individual neurons, we completed immunofluorescence analysis using an antibody for neuron-specific beta tubulin III, Tuj1. Syncytia structures were positively stained for Tuj1 (**Fig. 6E**), although the morphology was no longer consistent with a terminally differentiated neuron. We observed similar structures in the fetal cortex derived neurons (**Fig. 6E**). Taken together, these results demonstrate that infection by HCMV causes post-mitotic forebrain neurons to fuse and express markers of proliferation without signs of cell death.

### HCMV infection damages forebrain neuron function

Previously, our group demonstrated that infection negatively impacts calcium signaling in fibroblasts, NPCs, and mixed neural populations from cortical organoids by depleting calcium stores and compromising ATP/KCl receptor function (26). To assess whether this was also true in forebrain neurons, we used live ratiometric calcium imaging via FURA-2AM. When selecting for KCl-responsive cells (neurons), HCMV infected cells (eGFP+) demonstrated reduced calcium baselines compared to mock controls (**Fig. 7A and B**). This was consistent across all tested lines, regardless of genetic background. Further, infection reduced the average response to KCl stimulation within this neuronal subgroup, highlighting HCMV’s potential to reduce calcium mobilization in response to KCl stimulation (**Fig. 7C**).

**FIG 7.**
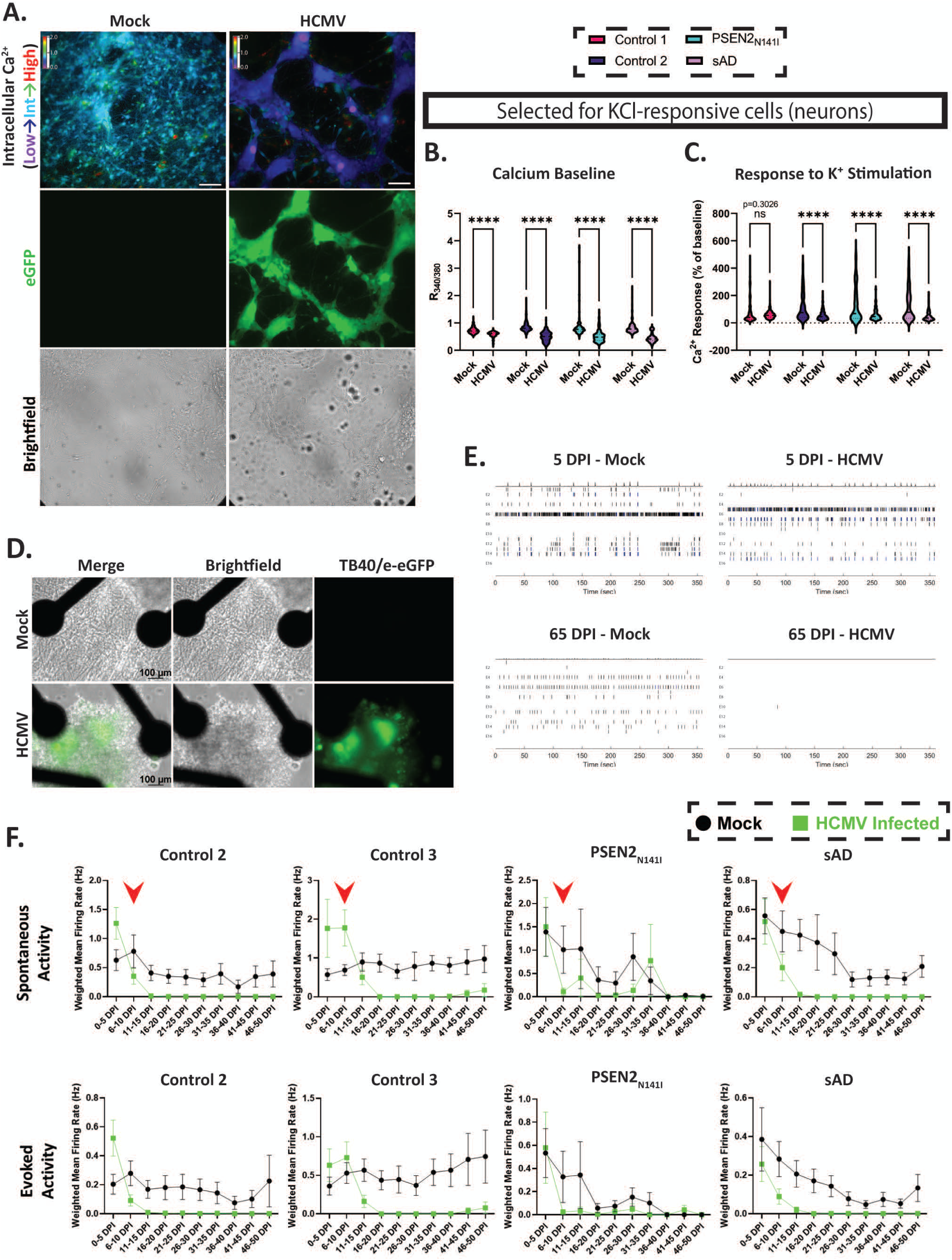
HCMV infection affects forebrain neuron function via dysregulated calcium dynamics. (A) Initial FURA-2AM imaging reveals widespread decreases in baseline calcium among infected forebrain neuron cultures (7 dpi). (B) Selecting for potassium chloride-responsive cells (neurons), HCMV’s effects on baseline calcium are significant across all tested lines (n = 188-296 cells per condition). (C) Upon stimulation with KCl, HCMV dampens calcium influx (n = 188-296 cells per condition). (D) Representative images of mock-treated and infected forebrain neurons plated onto multielectrode arrays (MEAs) at 65 dpi. (E) Representative raster plots showing electrical activity at 5 and 65 dpi. Spontaneous action potentials are nearly eliminated in HCMV cultures over time, whereas mock-treated cells still retain functional capabilities. (F) HCMV’s effects on action potential generation (both spontaneous and evoked) are present as early as 7 dpi, correlating with calcium imaging findings in A-C (timepoint indicated by red arrow). By 20 dpi, most electrical activity has ceased This is finding is ubiquitous across all tested lines. Analyses are restricted to MEA wells that demonstrate activity above threshold (wMFR > 0.15) (n = 5-15 wells per condition). Statistics in panels B and C were completed using 2-way ANOVA. *, p < 0.05; **, p < 0.01; ***, p < 0.001

Having established that HCMV can alter calcium signaling, we next tested the effect of infection on neuronal electrophysiological function. Cells were plated onto multielectrode arrays (MEAs) and cultured for 50 days (D74 of differentiation). During this period, cells were evaluated for neuronal maturation via action potential generation. Complementing our observations in calcium imaging, electrophysiological activity increased until D54 and plateaued, indicating functional maturity. Subsequently, each culture was infected with TB40/E-eGFP or mock treated, and recordings were conducted on each plate 3 times per week. Cultures were maintained in this paradigm for a minimum of 50 DPI (D124 of differentiation). Within 5 days, small punctate regions of eGFP were present within the MEA wells. As indicated by the 7 DPI TUNEL experiments (**Fig. 6A and B**), infection over long periods of time did not result in apparent cell death, although neuronal processes (axons, dendrites) disappeared by 65 dpi in infected cultures (**Fig. 7D**). Further, long-term HCMV infection effectively eliminated any spontaneous and evoked action potential generation within forebrain neuron cultures, as observed by Raster plot (**Fig. 7E**). Time-course analysis of spontaneous and evoked activity measured via weighted mean firing rate within cultures highlighted a rapid decline in action potential generation between 0 and 15dpi across all cell lines (**Fig. 7F**). The timeline of this decline coincided with the reductions in baseline calcium and KCl response observed via calcium imaging at 7 dpi (**Fig. 7A-C**, timepoint denoted by red arrows in **Fig. 7F**). Taken together, these data demonstrate that HCMV infection of mature forebrain neurons disrupts cell calcium signaling and impairs the electrophysiological neural function.

## DISCUSSION

HCMV infection is widely known to induce severe developmental neurological deficits due to the clear susceptibility of NPCs to HCMV based on both in vitro studies and in vivo evaluation of where HCMV is localized in the fetal brain (19, 20, 59, 60). However, there is also a potential link between HCMV seropositivity and age-related neurodegeneration in Alzheimer’s disease (61–63), which suggests that HCMV may also impact the health and function of terminally differentiated adult neurons. Here we show that HCMV robustly infects terminally differentiated iPSC-derived neurons and dramatically disrupts their structure and function.

To test if HCMV could infect terminally differentiated neurons, we differentiated iPSCs from both healthy individuals and patients with Alzheimer’s disease in vitro for 84-120 days prior to HCMV infection. Although we do recognize a limitation that iPSC-derived neurons lack full adult in vivo neuron characteristics (64–66), the neurons used here expressed clear markers of post-mitotic glutamatergic and GABAergic forebrain neurons and showed functionally mature calcium and electrophysiological properties (**Figs. 2, 7, S1A**). Action potential generation is a fundamental feature of mature neurons and results from coordinated activity of a variety of voltage gated ion channels (67). Immature iPSC-derived neurons will generally fire a single action potential with low amplitude, whereas mature iPSC-derived neurons will fire rapid chains of action potentials (67). In this regard, we observed iPSC-derived forebrain neurons from all five iPSC lines produced actional potential bursts indicative of neuronal maturation.

Previous studies have shown either a decrease in susceptibility to HCMV infection or a more limited impact of HCMV infection as NPCs were pushed toward a more differentiated state (19, 68). However, we did not observe this to be the case as neurons in culture for 84-120 days were robustly infected at an MOI of 3 (**Figs. 4A, 5D, 6D-E**, **7A**). The genetic background of the patient cell line also did not impact infection efficiency as all five iPSC lines showed similar results. Additionally, similar data were found in fetal cortex derived neurons indicating that these results were not simply a feature of the iPSC system. Together, these data demonstrate that HCMV infection is possible even when occurring at points substantially past terminal differentiation.

We and others have shown that NPCs undergo cell death in response to HCMV infection (19, 50), but we did not observe significant cell death in infected terminally differentiated neurons (**Fig 5A**). However, infected neurons lost characteristic morphological and functional features of neurons and showed a reactivation of proliferative markers. It is well known that HCMV over-rides the host cell cycle regulation in order to optimize viral gene expression, viral DNA replication, and viral production (69). In congenital HCMV infection, it is possible that some loss of NPCs is not detrimental to overall viral propagation because of the large NPC pool available in the early developing brain that could be used to maintain optimal viral replication. However, with a finite number of post-mitotic neurons in the adult brain, overt cell loss could impede viral spread particularly if viral replication and/or production is less efficient. It is not known why or how HCMV would have different effects in progenitor cells compared to terminally differentiated neurons, so future studies assessing transcriptional and proteomic profiles will be needed.

We and others have assessed the impact of HCMV on calcium function in NPCs (26) and other cell types (27, 70, 71), but the functional impact HCMV has on terminally differentiated neurons had not been previously established. Similar to results from NPCs, we found that HCMV dramatically reduced calcium baseline levels and impaired the calcium response to a KCl depolarizing stimulus (**Fig. 7A-C**), which is likely due to disruptions in voltage-gated ion channel function (26). Action potential generation was effectively eliminated within two weeks of infection (**Fig. 7F**). These data provide important insight into potential pathological effects of HCMV in the adult brain. For example, HCMV seropositivity is correlated with Alzheimer’s disease and a faster rate of cognitive decline (61). Action potentials drive communication between neurons in the CNS, and maintained neuronal connectivity is essential for cognitive health. Therefore, it is possible HCMV infection in adult neurons significantly reduces neuronal communication, thereby impacting overall cognitive function. More research is needed to directly assess HCMV infection on overall disease pathology, but the data presented here suggest that neuronal HCMV infection could induce significant functional impairment.

Upon observing infection in terminally differentiated neurons, one question we had was whether the infection progressed through full viral replication and if infected neurons could produce infectious virus. To answer the first question, we used the TB40/e-GFP+mCherry dual labeled virus in which mCherry is driven by the UL99-produced late viral protein pp28. Importantly, pp28 protein is only synthesized after the onset of viral replication (72). Using this viral construct, we observed a steady rise in pp28-driven mCherry expression in terminally differentiated neurons across all samples beginning around 100 hpi (**Fig. 4D**). Interestingly, we found mCherry expression in the neurites of infected cells (Fig. 6C) consistent with neuronal transport observed with other herpesviruses (73–75). This could indicate a potential cell-cell mediated mechanism for viral propagation in the adult brain to sites away from the initial infection. Moreover, we found robust mCherry expression in structures reminiscent of virion assembly compartments (**Fig. 6D**) surrounded by areas of organized multi-nucleated syncytia-like structures (**Fig D-E**). Syncytia formation is widely observed in HCMV infected tissues and is thought to be involved in viral spread to neighboring cells (58, 76, 77), so the structural changes associated with HCMV infection in terminally differentiated neurons may also play a role in viral propagation in the adult brain.

To assess whether infected neurons could produce infectious virus, we collected the conditioned medium and measured viral titers on naïve ARPE19 epithelial cells (**Fig. 4E**). Previous studies using NPC cultures have found that infected NPCs can propagate the infection to other cells in the culture (18), but others have found that infected NPCs, even infected at an MOI of 20, produced poorly infectious virions (68). We observed that iPSC-derived neurons from all five lines produced highly infectious virions (**Fig. 4E-F**), but we did note that neurons generated more non-functional virus that what is produced from fibroblasts (**Fig. 4F**). It is not clear why this is the case, but there may be differences in how HCMV is packaged and released by neurons compared to fibroblasts. Nevertheless, it will be interesting to further test whether the non-functional viral particles still have an impact on neuronal health and function through the release and expression of tegument proteins.

Together, these data show that terminally differentiated, functionally mature human neurons are robustly infected by HCMV, which results in a substantial decline in functional properties. Although the detrimental effect of congenital HCMV infection has been well established, the data presented here suggest that HCMV infection in terminally differentiated neurons is similarly detrimental and could impact overall brain health in the adult population.

## METHODS

### Cell Culture and Differentiation

Undifferentiated induced pluripotent stem cell (iPSC) lines originated from reprogrammed fibroblasts or patient blood cells (Control 1 – Coriell GM02183 [Fibroblast line reprogrammed to iPSCs]; Control 2 – Coriell GM03814 [Fibroblast line reprogrammed to iPSCs]; Control 3 – WiCell PENN022i-89-1 [iPSC]; PSEN2_N141I_ – Coriell AG25370 [iPSC]; sAD – Coriell AG27607 [iPSC]). Stem cells were maintained under feeder-free conditions on Geltrex-coated (Gibco) 6-well plates, with daily replacements of Essential 8 medium (Thermo Fisher Scientific). All iPSCs were grown for a minimum of three passages post-thaw prior to differentiation.

Patterned differentiation of iPSCs toward neural progenitor cells (NPCs) was conducted using dual-SMAD inhibition (STEMdiff SMADi Neural Induction Kit, STEMCELL Technologies). To promote complete and uniform generation of NPCs, cells were maintained in SMADi medium (with daily media changes) for a minimum of three passages prior to further differentiation to neurons. During each passaging step, NPC monolayers were dissociated to single-cell solutions using accutase solution (STEMCELL Technologies) and subsequently replated at a density of 2x10^5^ cells/cm^2^ onto Geltrex-coated 6-well plates. Cultures were maintained for 6 days between passages. At day 18 of differentiation, NPCs were again dissociated and plated at a density of 1.25x10^5^ cells/cm^2^.

NPC (iPSC-derived) patterning toward forebrain neuron fate was conducted using a commercially available kit (STEMdiff Forebrain Neuron Differentiation Kit, STEMCELL Technologies). Cells were maintained in forebrain neuron differentiation medium for 6 days before being dissociated into a single-cell solution and plated onto either Poly-L-Lysine- and Laminin-coated glassware/ Geltrex-coated plasticware at varying densities (6 well plate – 5.2x10^4^ cells/cm^2^; 24 well plate - 1.84x10^4^ cells/cm^2^; coverslip - 1.58-1.84x10^4^ cells/cm^2^; MEA – 1.36x10^3^ cells/cm^2^; 96 well plate – 1.88x10^5^ cells/cm^2^). Immature neuronal precursors were matured using STEMdiff Forebrain Neuron Maturation Medium (STEMCELL Technologies) until use.

Fetal-derived neuronal cultures were differentiated from neurospheres, as described by Ebert et al. (36). Briefly, 8-12 week post-conception cortical tissues (M045 and G010) were originally obtained from the University of Washington Birth Defects Research laboratory under an approved IRB protocol. Tissue samples were dissociated, cultured as floating aggregates, and passaged by mechanical chopping for ∼20 weeks in DMEM/F12 medium supplemented with 20ng/ml EGF, 20ng/ml FGF, and 2% B27. Neurospheres were aliquoted and frozen for later use. To induce differentiation, neurospheres were thawed, dissociated, and plated onto laminin coated coverslips in minimal Neurobasal medium supplemented with 2% B27. The in vitro use of human fetal tissue was approved by the Medical College of Wisconsin Institutional Review Board (PRO00025822).

### Viruses

HCMV viruses TB40/E-eGFP (37, 38) and dual fluorescently tagged TB40/E expressing IE2-2A-eGFP and UL99-mCherry, generously provided by Eain Murphy (SUNY Upstate Medical University, Syracuse, NY), originated from transfection of both a bacterial artificial chromosome (BAC) encoding each HCMV subvariant and a UL82-encoding plasmid into MRC-5 fibroblasts. This was accomplished using electroporation (260 mV for 30 ms; 4 mm-gap cuvette) via a Gene Pulser XCell system (Bio-Rad). Subsequently, generated stocks of each fibroblast-derived virus were used to infect ARPE-19 cells, generating epithelial-derived TB40/E-eGFP and TB40/E-eGFP/mCh. Upon collection, medium containing virus was processed by centrifugation (Sorvall WX-90 Ultracentrifuge and SureSpin 630 rotor; ThermoFisher Scientific), using a sorbitol cushion (20% sorbitol, 50 mM Tris-HCl [pH 7.2], 1 mM MgCl_2_) at 20,000 x g for 1 hour. Stock titers were determined by applying virus to ARPE19 cells within a 96 well plate and conducting a limiting dilution assay. At 2 weeks post-infection, GFP^+^ wells were identified, and infectious units were established in infectious units per milliliter (IU/mL). Infections were conducted at an MOI of 3 unless otherwise stated. Infections were conducted by applying virus-containing (or virus-free for mock) media to cells for 2 hours, with constant agitation. After two hours, media (+/-virus) was removed from each well, cells were washed 1x with Dulbecco’s Phosphate-buffered saline (PBS, Gibco), and fresh media was added.

### Viral titers and infectivity

Viral titers were assessed by collecting conditioned media from infected cultures at 14 DPI (iPSC-derived Forebrain Neurons), 7 DPI (fetal-derived neurons) and 4 DPI (neural progenitor cells). These timepoints varied due to culture viability. Serial dilutions of conditioned media were administered to plated ARPE-19 cells in a 12 well dish (in duplicate), and the cells were allowed to incubate for 1 week. After incubation, cells were stained using an antibody to viral protein immediate early 1 (IE1, added 1:500), and the IE1^+^ cells quantified using Goat α-mouse AF488 (1:1000). Final results were reported as infectious units per milliliter (IU/mL).

Using Qiagen’s DNeasy blood and tissue kit, DNA was isolated from the same neuronal/NPC conditioned media samples as used in the infectious unit assay. Isolated DNA was then assessed for the number of copies of viral immediate early gene UL123 using quantitative PCR (qPCR). Primer sets specific to UL123 were utilized (5′-GCCTTCCCTAAGACCACCAAT-3′ and 5′-ATTTTCTGGGCATAAGCCATAATC-3′). Subsequently, the UL123 copy number was divided by the infectious unit (IU) count obtained from the infectious unit assay. The resulting value addresses the infectivity of viral particles at 14 dpi.

### Protein and DNA analyses

Evaluation of relative viral-to-cellular DNA levels was completed using qPCR. Cells were collected for timepoints ranging from 1-5 days post-infection, with DNA from each being isolated via the DNeasy blood and tissue kit (Qiagen). Subsequently, primer sets for HCMV UL123 (5′-GCCTTCCCTAAGACCACCAAT-3′ and 5′-ATTTTCTGGGCATAAGCCATAATC-3′) and cellular GAPDH (5′-GTGGACCTGACCTGCCGTCT-3′ and 5′-GGAGGAGTGGGTGTCGCTGT-3′) (Integrated DNA Technologies) were used to determine relative viral DNA over the time course. 2x SYBR green PCR Master Mix (Bio-Rad, Thermo Fisher) was utilized for the qPCR. Data collection and analysis were completed using a Quantstudio 6 Flex real-time PCR machine (Thermo Fisher). Results utilized relative HCMV UL123 quantities being normalized to relative cellular GAPDH quantities.

Cells intended for protein analysis were plated at a density of 5x10^6^ per well of a Geltrex-coated (Thermo Fisher) 6-well plate. At 98 days of differentiation (14 DPI/mock treatment), cells were harvested and centrifuged to form a pellet. Pellets were frozen at -20C until use. Pellets were incubated in cold Triton-X lysis buffer (150 mM NaCl, 50 mM Tris-HCl [pH 8.0], 1 mM EDTA, 1% Triton X-100 [v:v], 1% protease inhibitor [v:v]) for 20 minutes and lysed via sonication. Sample concentrations were determined using a Pierce bicinchoninic acid (BCA) assay (Thermo Fisher). Using 30 µg of protein, samples were resolved by SDS-PAGE, using a 4-20% acrylamide gradient gel (Bio-Rad). Proteins were transferred to an Immobilon polyvinylidene difluoride (PVDF) membrane (Millipore) using a wet transfer Mini Trans-Blot Cell (Bio-Rad). Membranes were blocked for 1 hour at room temperature in Intercept (TBS) Blocking Buffer (LI-COR). Subsequently, membranes were incubated with primary antibody mix (Intercept Blocking Buffer, 0.2% Tween-20 [v:v], primary antibody) overnight at 4°C, with agitation. Membranes were washed three times using TBS-T (tris-buffered saline, 0.1% Tween-20) and secondary antibody solution (Intercept Blocking Buffer, 0.2% Tween-20 [v:v], 0.02% sodium dodecyl sulfate [SDS; v:v], secondary antibody) was applied for 30 minutes at room temperature. An Odyssey CLx fluorescent imaging system (LI-COR) was used to visualize protein banding. Primary antibodies used for western blotting in this study include rabbit anti-cyclin B1 (1:1000; Cell Signaling Technology), mouse anti-p21 (1:1000; Millipore-Sigma), and mouse anti-PCNA (1:500-600; Cell Signaling Technology). Secondary antibodies used in these experiments: donkey anti-rabbit 680RD (1:2000-3000; LI-COR) and donkey anti-mouse 800CW (1:2000-3000; LI-COR).

### Immunofluorescence and Live Imaging

iPSC- and fetal-derived neurons were plated onto Poly-L-Lysine/Laminin-coated (MilliporeSigma) coverslips at a density of 30-35,000 per coverslip. Cells were fixed via incubation in 4% paraformaldehyde for 15 minutes, washed with dPBS twice, and stored in fresh dPBS until use. Application of Triton X-100 solution (0.2% v:v, in PBS) for 10 minutes permeabilized cells. After washing once with PBS, blocking buffer (5% normal donkey serum [NDS], 5% normal goat serum [NGS], v:v, in PBS) was applied to the cells for 1 hour at room temperature. Coverslips were incubated in primary antibody buffer (primary antibody, 2.5% NGS, 2.5% NDS, 0.1% Triton X-100, in PBS) overnight at 4°C and secondary antibody solution (secondary antibody, 2.5% NGS, 2.5% NDS, 0.1% Triton X-100, in PBS) for 1 hour at room temperature. Nuclei were stained using Hoechst 33342 (1:1000 in PBS; Invitrogen). Coverslips were mounted on glass slides using Fluoromount-G Mounting Medium (Invitrogen) and sealed using clearcoat nail polish. Slides were imaged using a Zeiss LSM980 confocal microscope, with image analysis being conducted in Nikon Elements and Zeiss ZEN analysis software.

The following antibodies were used for immunofluorescence during this study: chicken anti-TUJ1 (1:250-300; GeneTex), rabbit anti-PAX6 (1:200-300; abcam), mouse anti-Ki67 (1:500; BD Biosciences), rabbit anti-VGLUT2 (1:500; Synaptic Systems), rabbit anti-GABA (1:100; Enzo), and chicken anti-vimentin (1:250-500; abcam). TUNEL staining (Click-iT TUNEL AlexaFluor 647; Invitrogen) was preformed according to the manufacturer’s specifications.

Live cell visualization of the virally encoded eGFP and mCherry fluorophores was completed using a Nikon TS100 inverted microscope. Timelapse live imaging was completed using an IncuCyte S3 in-incubator imaging system (Essen Biosciences). Brightfield, green fluorescent (ex. 440-480 nm; em. 504-544 nm), and red fluorescent (565-605 nm; em. 625-705) images were captured every two hours for seven days using a 10x objective. Incucyte 2022A software was used for background subtraction, image analysis, and video stitching.

### Calcium Imaging and Microelectrode Array

Immature neurons were plated onto Poly-L-Lysine/Laminin-coated (MilliporeSigma) coverslips at a density of 30,000 cells per coverslip. After D77 of differentiation, cells were infected with HCMV or mock-treated. At 7 DPI, cells were removed from neuronal maturation media, washed once with extracellular normal HEPES (ENH) solution (150 μM NaCl, 10 μM HEPES, 8 μM glucose, 5.6 μM KCl, 2 μM CaCl_2_, 1 μM MgCl_2_), and loaded with FURA-2AM calcium dye solution (2.5 µM FURA-2AM, 2% BSA [w:v], ENH) for 1 hour at room temperature. Coverslips were then washed again with ENH for 15 minutes before being mounted to a perfusion chamber. Prior to calcium recordings, both green fluorescent and brightfield images were obtained. To establish baseline readings, coverslips were superfused with ENH solution at a rate of 6mL per minute for 1 minute prior to initial stimulation. 10 µM ATP was applied to the cells for 1 minute to elicit a purinergic response, with a 1-minute ENH washout period immediately following. Then, KCl was administered to the cultures for 30 seconds to examine voltage-gated ion channel activity. Imaging ended with a 30 second washout period. A Nikon Eclipse Ti-U inverted microscope was used for imaging, and NIS-Elements Advanced Research (Nikon) was utilized for analysis and image processing. Within each field of view, 50 cells were selected for recording. This experiment was repeated using three regions per coverslip, across two coverslips, per condition. Plotted R_340/380_ ratios represent bound (340 nm) to unbound (380) calcium at baseline.

Cells were plated onto a PLL/Laminin-coated (MilliporeSigma) Cytoview microelectrode (MEA) plate (Axion Biosystems) at a density of 30,000 per well. Cells were recorded three times weekly using a Maestro MEA system (Axion Biosystems), analyzing both spontaneous and electrically evoked action potentials. For spontaneous recordings, cells were recorded for a six-minute window with the default settings for spontaneous recordings set within the AxIS software suite (Axion Biosystems). Analysis of spontaneous data was conducted using the Neural Metric Tool (NMT, Axion Biosystems). In NMT, the criterion for an “active electrode” was defined as 5 spikes/minute, and coincidence artifacts were removed. Additionally, in generating time course data, wells that never demonstrated activity were removed from any subsequent analysis. Data are reported as weighted mean firing rate (wMFR). Electrically evoked potentials were recorded over a duration of 2 minutes (following spontaneous recordings), with 0.5 V stimulations (400 µs duration, each) occurring every 10 seconds. As with spontaneous recordings, evoked recordings were processed using NMT to determine active electrodes and remove coincidence artifacts. Additionally, any spikes occurring within 2 ms of the electrical stimulation were discarded to ensure detected action potentials were not artifactual. Neuron-free wells produced evoked wMFR values of around 0.15 Hz, leading to a background subtraction of 0.15 Hz from all wMFR values generated by NMT (evoked data only). As with spontaneous activity data, wells never exceeding this 0.15 Hz threshold were removed from further analyses.

### Statistical Analysis

All statistical testing was completed using the GraphPad Prism software suite. Utilized statistical tests are indicated in figure legends, with significance being defined as * < 0.05 (** < 0.01, *** < 0.001, **** < 0.0001).

## ACKNOWLEDGMENTS

We would like to thank Dr. Tom Shenk for access to the HCMV IE1 antibody and Dr. Eain Murphy for the fluorescently labeled viruses. Further, we appreciate the help of members of the Ebert and Terhune labs, for their helpful advice in the preparation of this manuscript.

Research reported in this publication was supported by the National Institute of Allergy and Infectious Diseases division of the National Institutes of Health under award number R01AI132414 (S.S.T. and A.D.E.). The content is solely the responsibility of the authors and does not necessarily represent the official views of the National Institutes of Health. These studies have also been supported by a generous philanthropic gift by The Stead Family Foundation to define the impact of infection and inflammation on brain health.

## FIGURE LEGENDS

**FIG S1 iPSC-derived and fetal-derived neuronal cultures are representative of cell types in the human forebrain.** (A) Both excitatory, glutamatergic (vGLUT2^+^) and inhibitory, GABAergic (GABA^+^) neuronal subtypes were present in the iPSC-derived culture system at D84 of differentiation. Additionally, limited populations of Vimentin+ astrocytes and Ki67+ proliferative cells (neural progenitors) were also observed. (B) Similar neuronal populations were found in fetal progenitor-derived cultures.

**FIG S2 Infection of forebrain neurons with HCMV is possible over a range of MOIs.** (A) At 9 dpi, forebrain neurons demonstrate markers of viral immediate early (IE2) and late (UL99) gene synthesis at all tested MOIs (0.5, 1, 3 IU/cell). An MOI of 3 produced the most fluorescent signal. (B) At 15 dpi, infected cultures continue to demonstrate IE2/eGFP and pp28-mCherry signal, though differences in intensity disappear.

**FIG S3 Fetal-derived neurons demonstrate HCMV spread**. Infection of fetal-derived neurons with TB40/e-GFP reveals progressive increases in fluorescent signal starting as early as 4 dpi. Large eGFP^+^ plaques are visible by 10 dpi, accompanied by a significant alteration to cell morphology. Scale bars = 200 µm.

**VID S1 TB40/e infection drives syncytia formation in forebrain neuron cultures**. 3D render of syncytial structure using z stacks of 0.430 µm thickness. Blue = Hoechst 33342; Green = viral eGFP; Red = TUJ1; Magenta = Ki67.

