## Supplementary figures and images for "Human Cytomegalovirus Induces Significant Structural and Functional Changes in Terminally Differentiated Human Cortical Neurons"

### supplemental figures

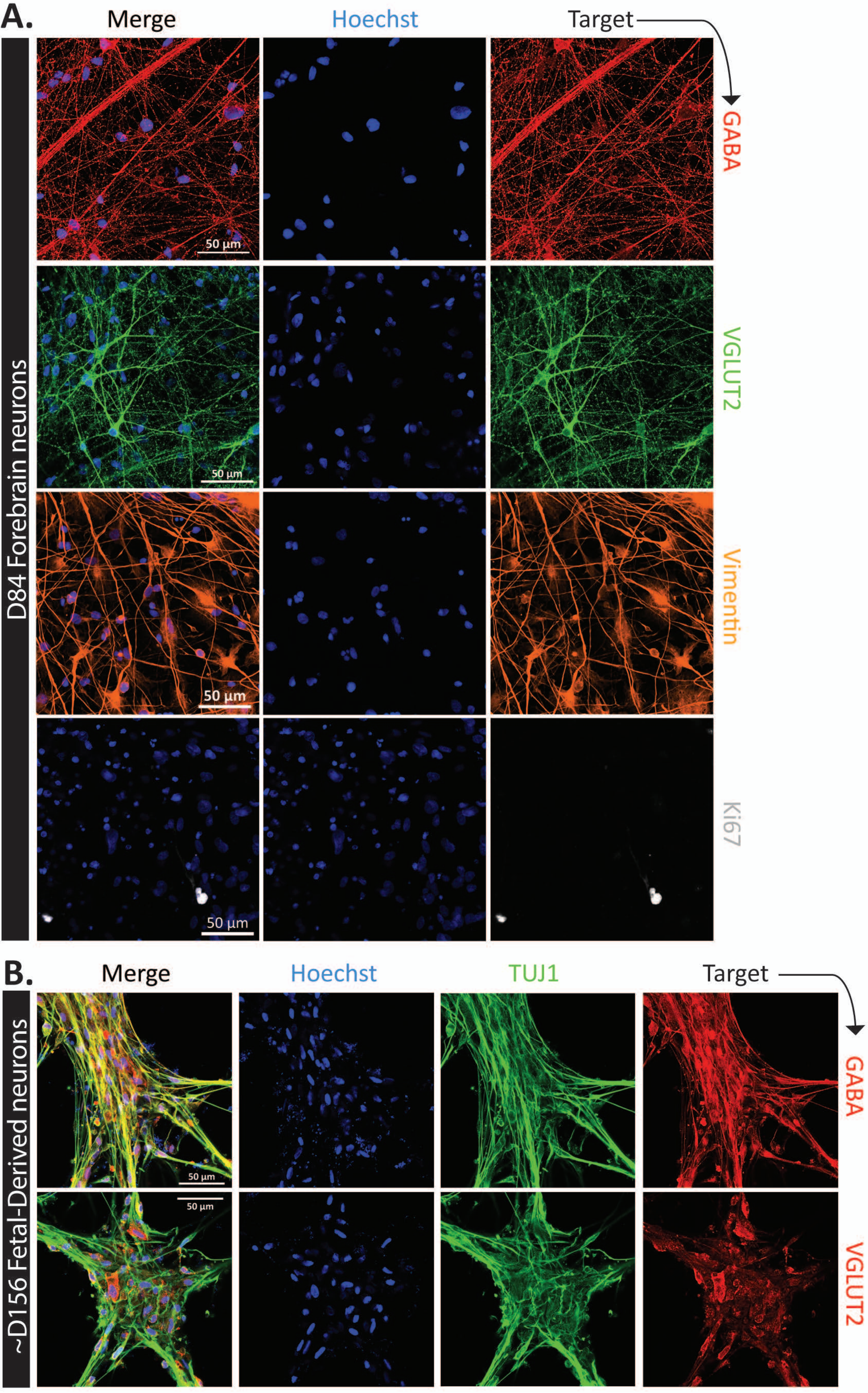

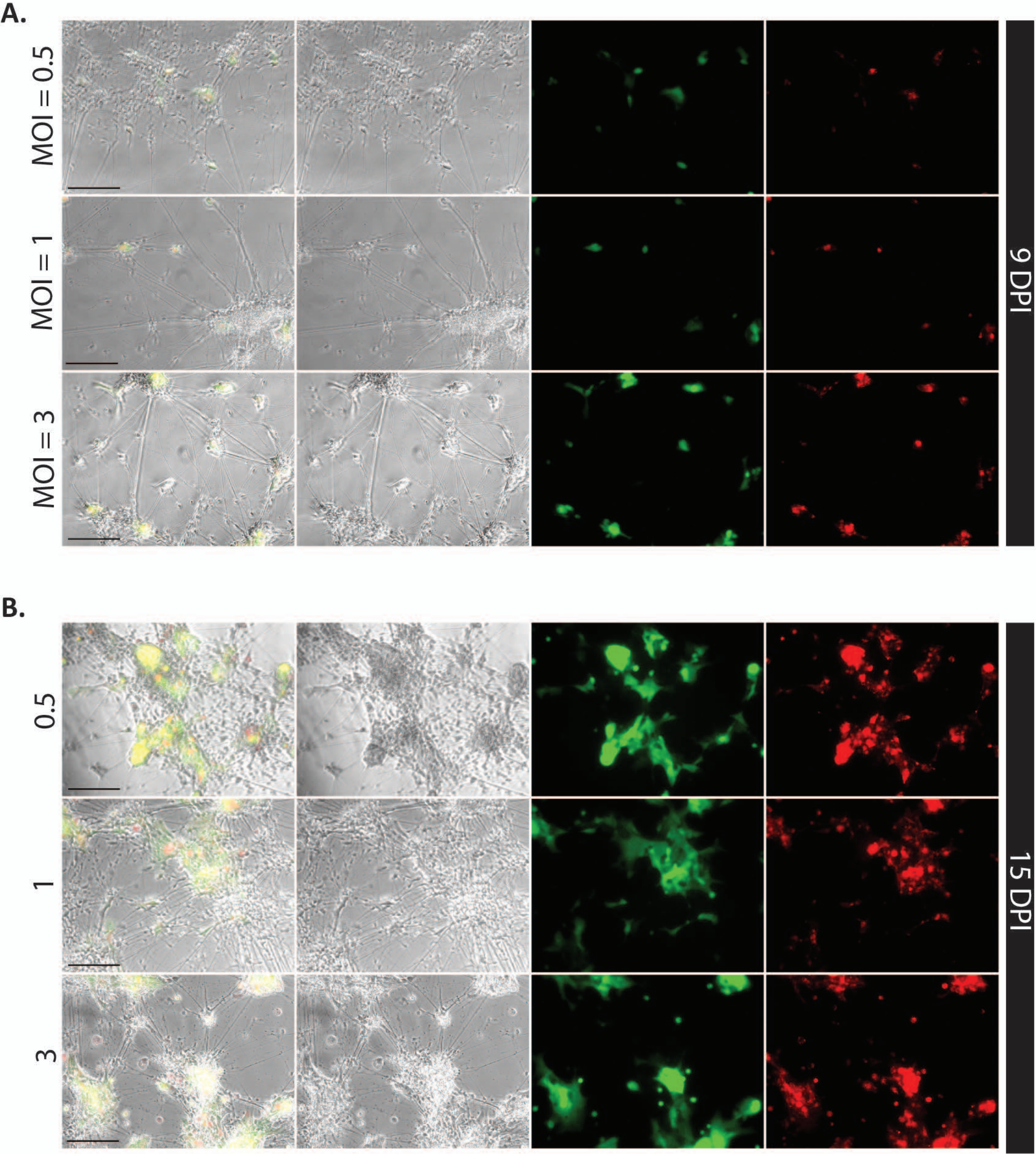

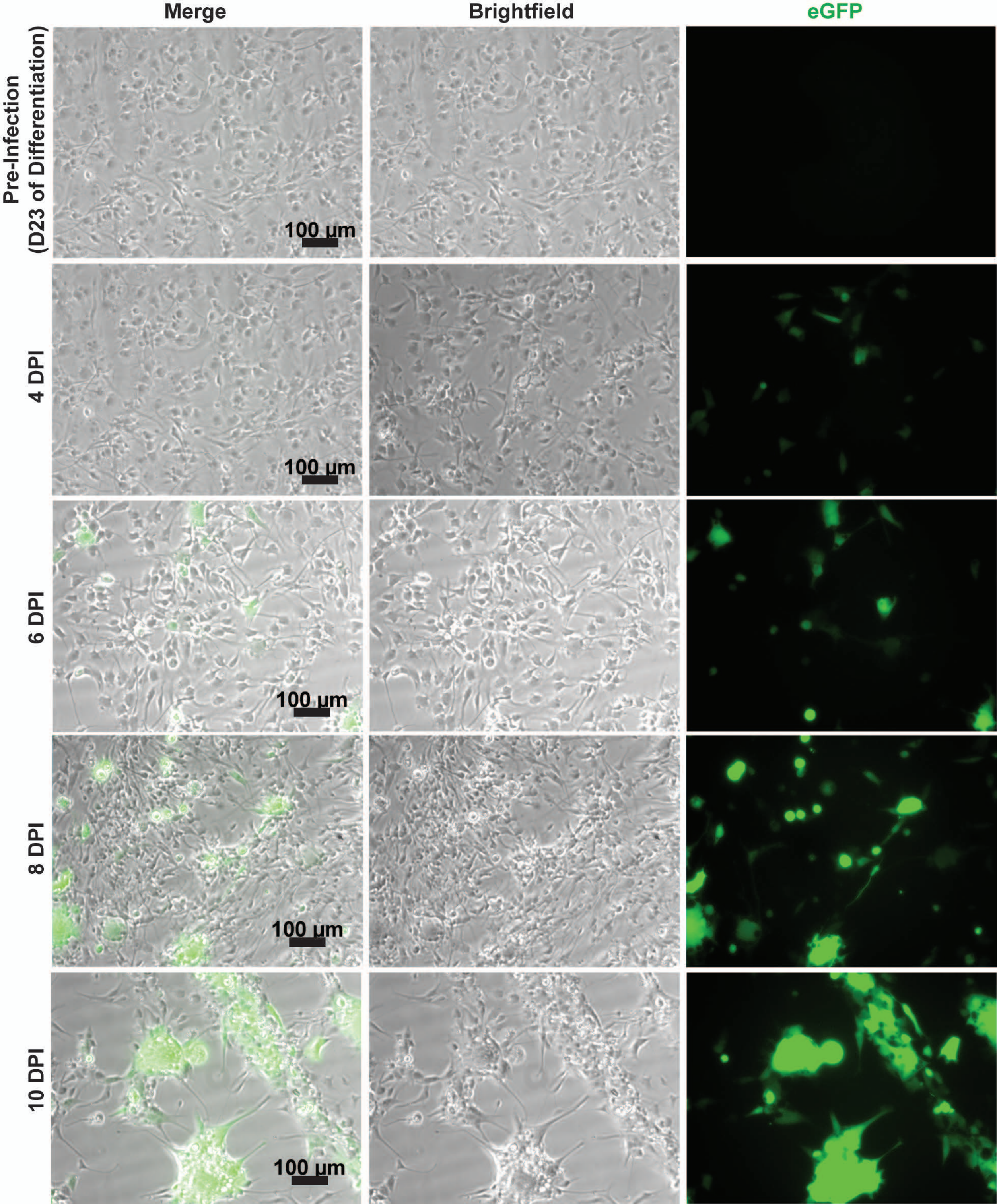
